# IReNA: integrated regulatory network analysis of single-cell transcriptomes and chromatin accessibility profiles

**DOI:** 10.1101/2021.11.22.469628

**Authors:** Junyao Jiang, Pin Lyu, Jinlian Li, Sunan Huang, Seth Blackshaw, Jiang Qian, Jie Wang

## Abstract

Although single-cell RNA sequencing (scRNA-seq) and single-cell assay for transposase accessible chromatin using sequencing (scATAC-seq) have been widely used, few methods can reliably integrate these data to perform regulatory network analysis. Here, we developed IReNA (Integrated Regulatory Network Analysis) for network inference through integrated analysis of scRNA-seq and scATAC-seq data, network modularization, transcription factor enrichment, and construction of simplified intermodular regulatory networks. Using public datasets, we showed that integrated network analysis of scRNA-seq and scATAC-seq data using IReNA outperformed currently available methods in identifying known regulators. IReNA facilitates the systems-level understanding of biological regulatory mechanisms, and is available at https://github.com/jiang-junyao/IReNA.

## Background

Dynamic changes of *trans*-regulators (e.g., transcription factors) and *cis*-regulatory elements (e.g., promoters) control gene expression in biological systems [1]. This fact makes it possible to infer gene regulatory networks using transcriptomic and epigenomic profiles. Recent advances in single-cell sequencing technologies provide new opportunities to reconstruct cell type-specific regulatory networks [2]. Single-cell transcriptomes have been widely detected through single-cell RNA sequencing (scRNA-seq). Currently, dozens of methods have used scRNA-seq data to infer regulatory networks, including top performing methods GENIE3 and PIDC [3–5]. Complementary to scRNA-seq, epigenomic profiling technique assay for transposase accessible chromatin using sequencing (ATAC-seq) measures accessible states of *cis*-regulatory elements to *trans*-regulators facilitating regulatory network inference [6]. Using scRNA-seq and bulk ATAC-seq data, we have developed a method to infer regulatory networks controlling retinal regeneration [7]. However, with the advent of the latest technology single-cell ATAC-seq (scATAC-seq), a new method for network inference is needed to integrate scRNA-seq and scATAC-seq data.

Besides network inference, dissecting regulatory networks to detect modules and to identify key regulators is another major challenge in systems biology. Clustering and decomposition are two frequently used methods for module detection [8]. Weighted correlation network analysis (WGCNA) performs hierarchical clustering of expression profiles to detect gene modules after correlation-based network inference [9]. Using single-cell transcriptomes, the SCENIC software combined GENIE3 and Rcistarget separately for network inference and identification of key transcription factors [10]. Although current methods can infer regulatory networks to identify gene modules and key individual regulatory genes, they do not construct a simple and statistically robust regulatory network among modules to provide new biological insights.

Here, we developed IReNA to perform regulatory network analysis through integrating scRNA-seq and scATAC-seq data. Since only scRNA-seq data are available for many biological samples, IReNA can also provide network analysis for using only scRNA-seq data. Using IReNA, we analyzed published single-cell RNA-seq and ATAC-seq profiles, reconstructed regulatory networks, identified key regulators, and revealed simplified regulatory networks among modules. In comparison with Rcistarget, one of the most frequently used existing methods, IReNA shows better overall performance at identifying known regulators.

## Result

### The framework of IReNA

To perform regulatory network analysis using single-cell RNA-seq data or integrating with ATAC-seq data, we developed IReNA which consists of two components: network inference and network decoding (Figure 1 and Figure S1). Two pipelines of network inference were developed separately for analyzing scRNA-seq data alone or integrating scRNA-seq data with bulk or single-cell ATAC-seq data. Using scRNA-seq data, potential regulatory relationships were firstly inferred by the tree-based ensemble method GENIE3. Then, transcription factor binding motifs were used to refine regulatory relationships. If bulk or single-cell ATAC-seq data is available, both transcription factor binding motifs and footprints were used to refine regulatory relationships. For scATAC-seq data, peaks were linked to genes and used to identify transcription factor binding motifs and footprints. Network decoding in IReNA includes network modularization, identification of enriched transcription factors and a unique function for the construction of simplified regulatory networks among modules.

**Figure 1.**
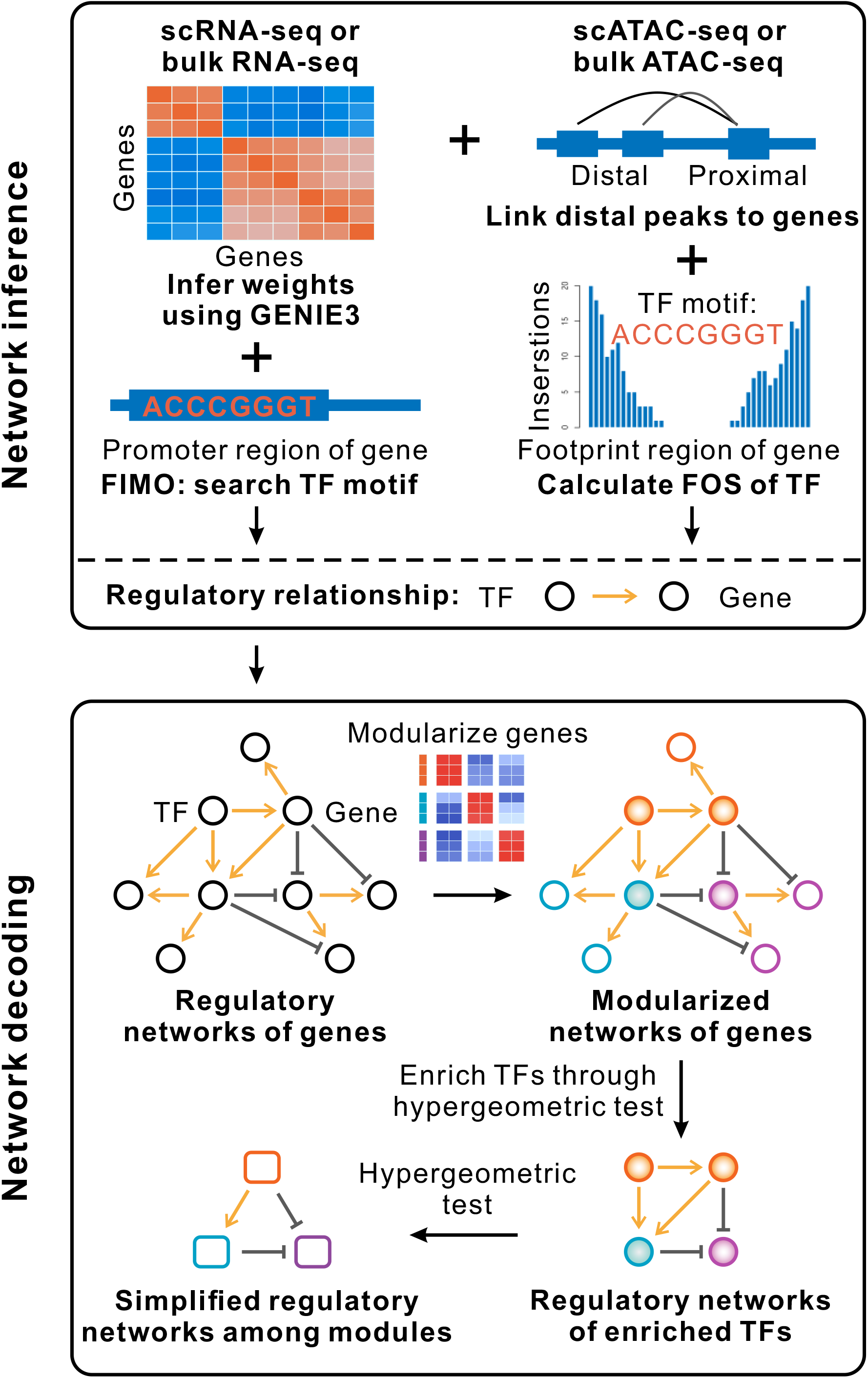
Flowchart of IReNA. IReNA consists of two components: network inference and network decoding. Network inference is performed according to weights calculated by GENIE3. FIMO is then used to identify binding motifs of transcription factors and refine regulatory relationships in cases where only gene expression profiles (single-cell or bulk RNA-seq data) are used. If bulk or single-cell ATAC-seq data is available, binding motifs and footprint occupancy score (FOS) are used to refine regulatory relationships inferred from expression analysis. Especially for single-cell ATAC-seq data, ArchR is used to link peaks and genes. Next, inferred networks are modularized and used to identify enriched transcription factors (TFs). Regulatory networks of enriched TFs are further decoded by establishing simplified regulatory networks among modules.

To illustrate the features and application of IReNA, we used public scRNA-seq and scATAC-seq data of hepatocytes from a mouse model of hepatectomy in the study of liver regeneration [11]. Although scRNA-seq and scATAC-seq data were obtained, the original study didn’t perform regulatory network analysis.

### Network analysis only using scRNA-seq data

Here, IReNA was applied to infer regulatory networks using scRNA-seq data alone. We firstly analyzed scRNA-seq data to identify genes used for network inference. Single-cell expression profiles of 2815 hepatocytes from the control and 48h liver tissues after hepatectomy were used to construct the trajectory of hepatocytes (Figure 2A). In the trajectory, we observed three branches, including the rest, activation and proliferation. Based on the trajectory, we calculated the pseudotime and identified 4014 differentially expressed genes (DEGs) including 165 transcription factors along the pseudotime. Meanwhile, we identified 45 transcription factors which are expressed in more than 5% hepatocytes but not statistically differential during pseudotime. Next, all 4059 DEGs and expressed transcription factors were modularized into five modules through K-means clustering of the smoothed expression profiles (Figure 2B). Genes in each module showed specific expression profiles. For instance, genes in the first and fifth modules were specifically expressed in the rest and proliferating hepatocytes, respectively. Function enrichment analysis revealed relevant biological functions enriched in each module of genes, including fatty acid metabolism, organic acid catabolism, cytoplasmic translation, autophagy and cell cycle (Figure 2C).

**Figure 2.**
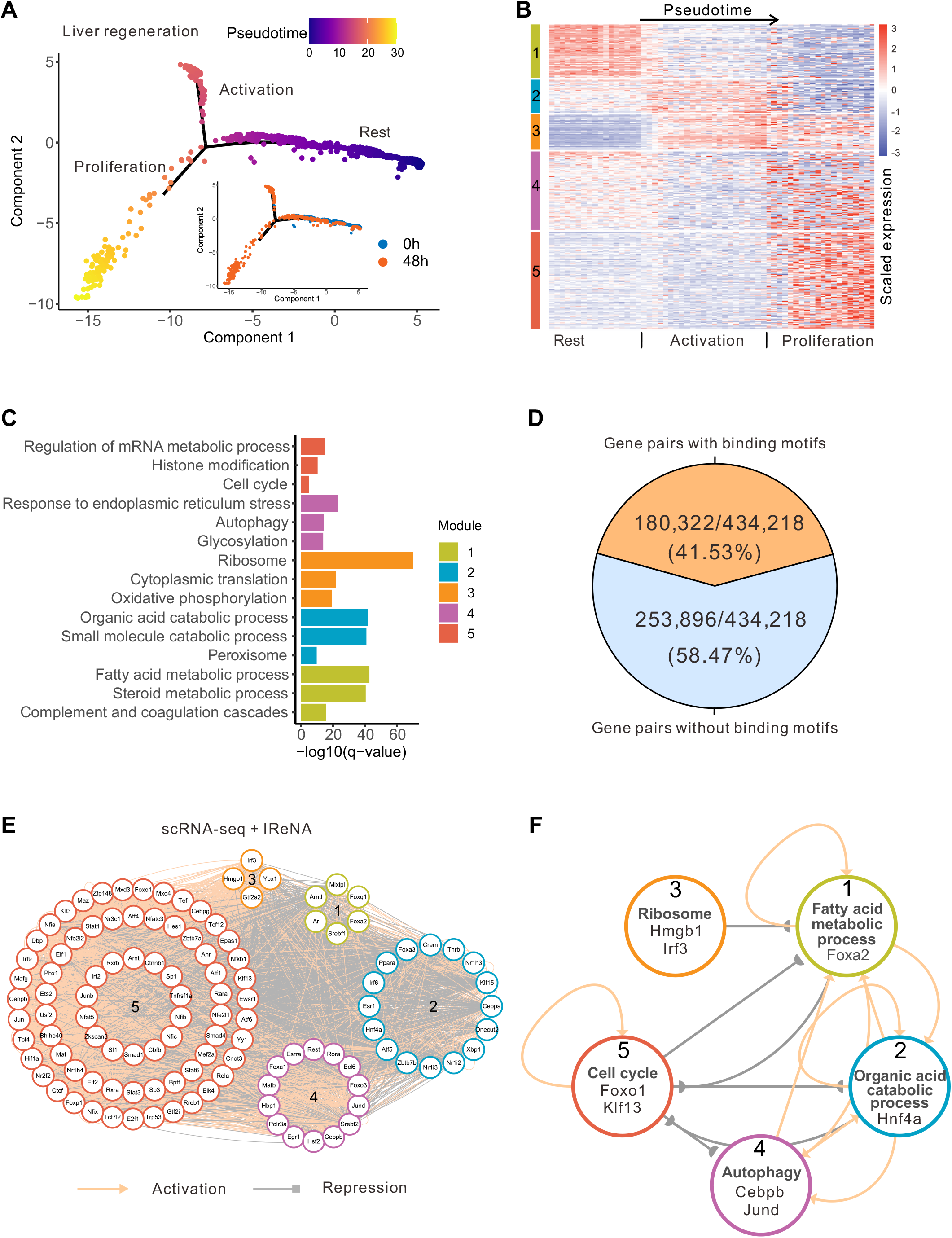
Regulatory network analysis of hepatocytes from liver regeneration. (A) Trajectory of 2815 hepatocytes from liver tissues at 0 and 48 hours after partial hepatectomy. Two trajectories are separately colored by pseudotime and samples. (B) Heatmap of scRNA-seq expression profiles of 4059 differentially expressed genes (DEGs) and expressed transcription factors. K-means clustering was used to divide genes into five modules. Rows represent genes, and columns indicate intervals of pseudotime separated by three branches. (C) Enriched functions of five modules of genes. (D) Fraction of regulatory gene pairs inferred by GENIE3 which contain transcription factor binding motifs in the promoter regions of target genes. (E) Regulatory networks of 41 enriched transcription factors obtained from analyzing scRNA-seq data alone. Color of each circle indicates the module. Grey edge represents negative regulation, and the yellow edge represents positive regulation. (F) Simplified regulatory networks among five modules obtained from analyzing scRNA-seq data alone. Representative biological functions and transcription factors were selected from Figure 2C and 2E to label each module.

We then applied GENIE3 to infer regulatory relationships of all DEGs and expressed transcription factors in hepatocytes. We identified 434,218 potential regulatory relationships, each of which has > 0.0001 weight and contains at least one transcription factor. For each regulatory relationship, Pearson’s correlation was calculated to determine the positive or negative regulation. To refine 434,218 potential regulatory relationships, we further analyzed transcription factor binding motifs in the promoter regions of genes. Totally, 180,322 regulatory relationships with binding motifs were used to reconstruct regulatory networks (Figure 2D). Meanwhile, regulatory networks were modularized according to five modules of genes identified through K-means clustering in Figure 2B. Statistically analyzing modular regulatory networks, we identified 115 transcription factors which significantly regulated each module of genes (Table S1). We reconstructed regulatory networks of these 115 enriched transcription factors, which were also divided into five modules (Figure 2E).

To obtain a simple regulatory network among modules for providing a clearer picture of biological regulatory relationships, we statistically analyzed modular regulatory networks of enriched transcription factors. Fifteen significant regulatory relationships among modules were identified and used to establish simplified regulatory networks among modules (Figure 2F). In simplified intermodular regulatory networks, we observed that transcription factors related to fatty acid metabolism, organic acid catabolism and autophagy significantly activated each other. In return, these factors significantly repressed transcription factors controlling cell cycle regulation. These indicate that inhibition of transcription factors related to fatty acid metabolism, organic acid catabolism and autophagy would activate cell cycle progression. Transcription factors relevant to ribosome repressed transcription factors of fatty acid metabolism, suggesting transcription factors regulating ribosome have synergistic effects on regulators of cell cycle.

### Network analysis through integrating scRNA-seq and scATAC-seq data

Next, IReNA was used to infer regulatory networks through integrating scRNA-seq and scATAC-seq data from the study of liver regeneration. We analyzed scATAC-seq data of 7004 hepatocytes after hepatectomy using ArchR [12]. We identified 94,595 significant peak-to-gene links, each of which had high peak-to-gene correlation, e.g., *Rora* and *Mlx* (Figure 3A). We further uncovered transcription factors binding to peaks, and identified 386,597 regulatory relationships of transcription factors to genes. Among 386,597 regulatory relationships, 154,601 regulatory relationships had high footprint occupancy scores. Overlapping 154,601 regulatory relationships with 434,218 potential regulatory relationships inferred by GENIE3, we refined 47,721 regulatory relationships to reconstruct regulatory networks which consisted of 3185 genes.

**Figure 3.**
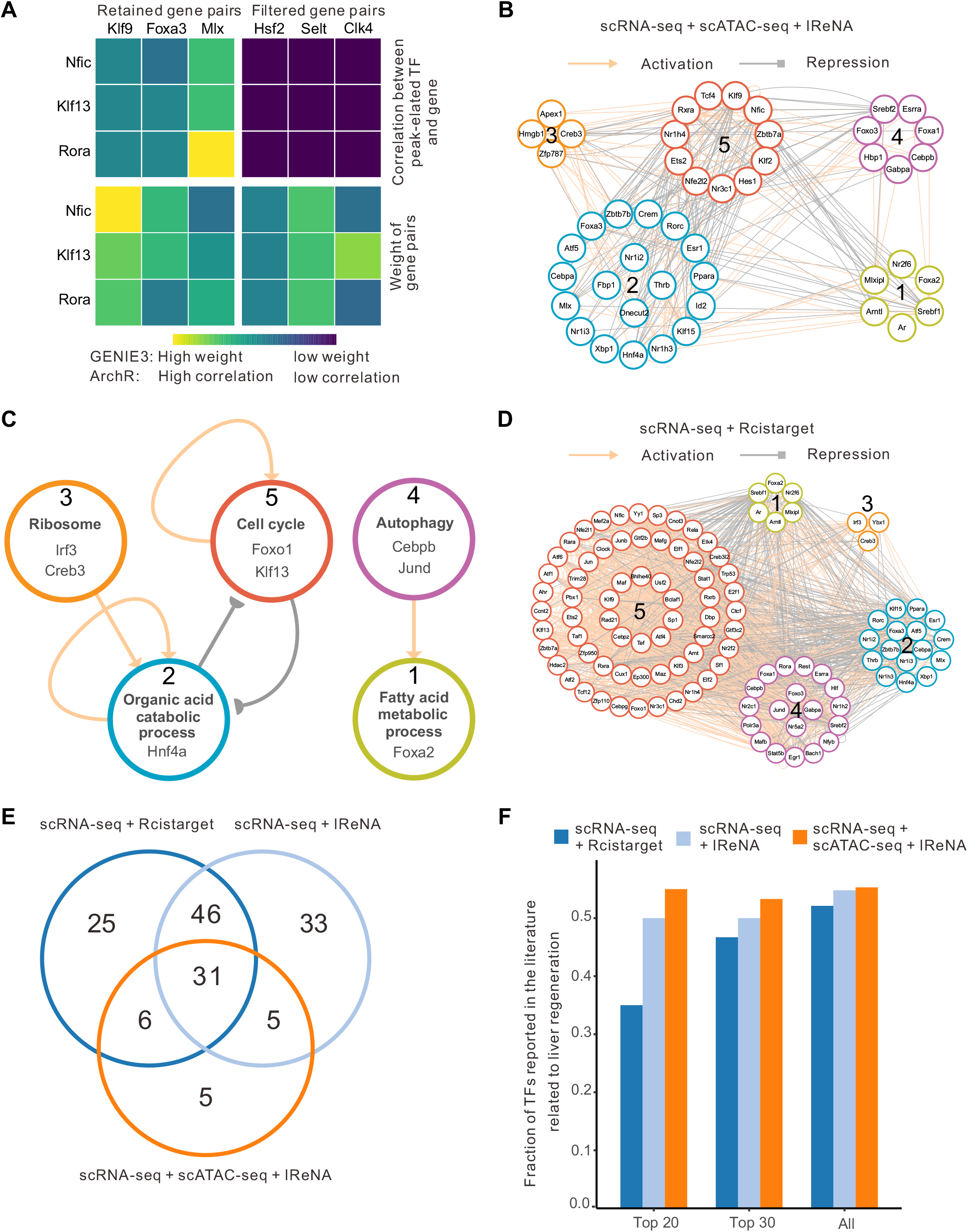
Comparison of regulatory networks related to liver regeneration. (A) Heatmap of peak-to-gene correlations (top panel) calculated by ArchR and weights of gene pairs (bottom panel) calculated by GENIE3. In the top panel, the gene in the column represents a transcription factor which has the binding motif in the peak. Gene pairs with both high correlations and high weights are used to infer regulatory networks of integrating scRNA-seq and scATAC-seq data. (B) Regulatory networks of 47 enriched transcription factors obtained through the integrated analysis of scRNA-seq and scATAC-seq data. Color of the circle indicates the module. Grey edge represents negative regulation, and the yellow edge represents positive regulation. (C) Simplified regulatory networks among modules obtained through the integrated analysis of scRNA-seq and scATAC-seq data. (D) Regulatory networks for 108 enriched transcription factors from scRNA-seq data analysis using Rcistarget. (E) Venn diagram of enriched transcription factors from three types of regulatory networks, one from the integrated analysis of scRNA-seq and ATAC-seq data using IReNA (ATAC-seq + scRNA-seq + IReNA), one from scRNA-seq data analysis using IReNA (scRNA-seq + IReNA) and one from scRNA-seq data analysis using Rcistarget (scRNA-seq + Rcistarget). (F) Fraction of enriched transcription factors reported in the literature related to tissue regeneration. Top 20, top 30 and all enriched transcription factors were compared for all three types of regulatory networks. Enriched transcription factors were ranked according to FDR value or normalized enrichment score.

Analyzing modular regulatory networks of 3185 genes, we identified 47 transcription factors which significantly regulate gene expression in each module. We then reconstructed modular regulatory networks of 47 enriched transcription factors (Figure 3B). Six significant regulation relationships among modules were identified to reconstruct simplified regulatory networks among modules (FDR < 0.05, Figure 3C). We observed that intermodular regulatory networks from the integrated analysis of scRNA-seq and scATAC-seq data are consistent with intermodular regulatory networks obtained from analyzing scRNA-seq data alone. In liver regeneration, the module of transcription factors related to organic acid catabolism of hepatocytes, e.g., *Hnf4a*, significantly repressed cell cycle. This suggests that the inhibition of hepatocyte metabolism may promote liver regeneration.

### Performance of IReNA on identifying reported transcription factors

To directly compare IReNA with other methods on network analysis, we analyzed scRNA-seq data to reconstruct regulatory networks using the Rcistarget package from the SCENIC software [10]. Using Rcistarget, we identified 108 transcription factors whose binding motifs were overrepresented in the promoter regions of 4059 DEGs and expressed transcription factors identified by scRNA-seq data analysis. We obtained 1716 significant regulatory relationships for 108 enriched transcription factors, and then reconstructed modular regulatory networks of enriched transcription factors (Figure 3D).

We then compared regulatory networks of enriched transcription factors inferred through three different approaches described above. Among 47 transcription factors identified by IReNA using integrated analysis of scRNA-seq and scATAC-seq data, 36 (76.60%) transcription factors were also present in regulatory networks inferred using IReNA analysis of scRNA-seq data alone (Figure 3E). By comparing gene regulatory networks obtained by analyzing only scRNA-seq data separately using IReNA and Rcistarget, we observed an overlap of 66.96% (77 in 115) of all enriched transcription factors. These indicated that a large fraction of transcription factors was identified by two methods.

To assess the significance of transcription factors identified using IReNA or Rcistarget, we manually examined whether these factors had been previously reported in the literature related to tissue regeneration. To compare different methods, we ranked the enriched transcription factors according to the significance in statistics (FDR for IReNA or normalized enrichment score for Rcistarget). We found that regulatory networks from the integrated analysis of scRNA-seq and ATAC-seq data have the best performance, 55.00% in top 20, 53.33% in top 30 and 55.32% in all enriched transcription factors were separately reported in regeneration-related literature (Figure 3F and Table S1). In regulatory networks inferred from analyzing scRNA-seq data alone, 50.00% in top 20, 50.00% in top 30 and 53.91% in all transcription factors were reported in regeneration-related literature. In regulatory networks inferred by Rcistarget, there were 35.00% in top 20, 46.67% in top 30 and 51.85% in all transcription factors reported in regeneration-related literature. Among three types of regulatory networks, the highest fraction of transcription factors was reported for regulatory networks from the integrated analysis of scRNA-seq and scATAC-seq data. These results indicate that integrated network analysis of scRNA-seq and scATAC-seq data using IReNA improved the precision of identifying known transcription factors. Moreover, IReNA shows better performance at identifying known regulators than the Rcistarget method.

To further demonstrate the performance of IReNA, we performed regulatory network analysis on another two datasets from heart regeneration and NASH. Prior to inferring gene regulatory networks controlling heart regeneration, we reconstructed the trajectory of 4884 cardiomyocytes from neonatal heart tissues (Figure S2A). We found that cardiomyocytes formed two distinct branches (named the activation branch and the proliferative branch) following myocardial infarction. We further identified and divided 4340 DEGs and expressed transcription factors into 4 modules (Figure S2B). Genes in the activation branch (module 2 and 3) were related to oxidative phosphorylation and muscle cell differentiation, whereas genes in the proliferative branch (module 4) are enriched for cell cycle (Figure S2C). Then, regulatory networks were inferred using the same three methods described above (Figure 4A-4B and Figure S2D-S2E). In comparison with regulatory networks inferred by Rcistarget, regulatory networks reconstructed by IReNA contained a higher fraction of transcription factors previously reported in the literature related to regeneration (Figure 4C and Table S1). The precision of identifying known regulators among top 20 enriched transcription factors were 80.00%, 70.00% and 45.00% separately for scRNA-seq + scATAC-seq + IReNA, scRNA-seq + IReNA and scRNA-seq + Rcistarget. We also observed that scRNA-seq + scATAC-seq + IReNA identified the highest fraction of known regulators for both top 30 and all enriched transcription factors.

**Figure 4.**
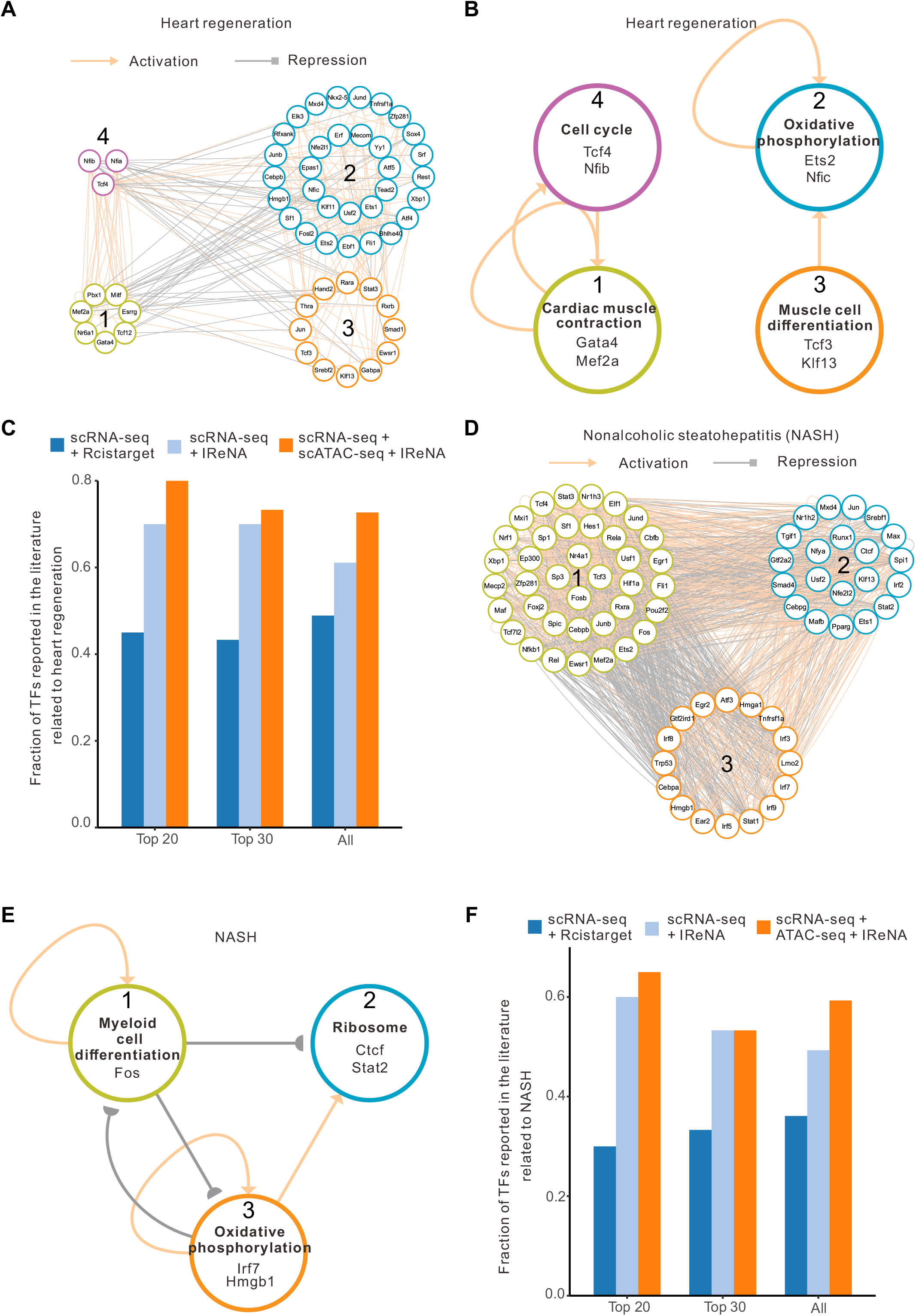
Regulatory network analysis for two studies of heart regeneration and NASH. (A) Regulatory networks of 54 enriched transcription factors obtained by integrating scRNA-seq and scATAC-seq data from heart regeneration. Modules are represented by the color of the circles. Grey edge represents negative regulation, and the yellow edge represents positive regulation. (B) Simplified regulatory networks among modules obtained by integrating of scRNA-seq and scATAC-seq data from heart regeneration. (C) Fraction of enriched transcription factors reported in the literature related to tissue regeneration. (D) Regulatory networks of 75 enriched transcription factors obtained by integrating scRNA-seq and bulk ATAC-seq data from nonalcoholic steatohepatitis (NASH). (E) Simplified regulatory networks among modules obtained by integrating scRNA-seq and ATAC-seq data from NASH. (F) Fraction of transcription factors reported in the literature related to NASH.

For the study of NASH, we used 2748 Kupffer cells to construct the trajectory (Figure S3A). 2742 DEGs and expressed transcription factors were identified and divided into three modules, which were separately enriched for myeloid cell differentiation, ribosome and oxidative phosphorylation (Figure S3B-S3C). In NASH, only bulk ATAC-seq data was available and used to refine regulatory relationships inferred from scRNA-seq data analysis (Figure S3D). We reconstructed three types of regulatory networks and a simplified regulatory network among modules (Figure 4D-4E and Figure S3E-S3F). Regulatory network comparison of NASH study has a similar trend with studies in liver regeneration and heart regeneration. IReNA analysis using scRNA-seq and bulk ATAC-seq data identified the most transcription factors (65.00% in top 20, 53.33% in top 30 and 59.26% in all enriched transcription factors) which are reported to associate with NASH, followed by IReNA analysis using scRNA-seq data alone (60.00% in top 20, 53.33% in top 30 and 49.33% in all), then is Rcistarget analysis using scRNA-seq data alone (30.00% in top 20, 33.33% in top 30 and 36.36% in all) (Figure 4F and Table S1). The integrated analysis of single-cell or bulk ATAC-seq data with scRNA-seq data overall substantially improved the reconstruction of gene regulatory networks, and had higher precision of identifying known transcription factors. In addition, regulatory networks reconstructed from scRNA-seq data alone using IReNA showed improved accuracy relative to those identified using Rcistarget.

## Discussion

In the study, we developed IReNA to perform regulatory network analysis including network inference and network decoding. In IReNA, gene regulatory networks are inferred through analyzing either scRNA-seq data alone or integrating scRNA-seq and scATAC-seq data. The regulatory relationships between transcription factors and target genes are firstly inferred according to the weights calculated by GENIE3 using scRNA-seq data. Then, transcription factor binding motifs are identified to refine regulatory relationships if only scRNA-seq data are available. When both scRNA-seq and ATAC-seq data are used, transcription factor binding motifs and footprints are applied to further refine transcriptional regulatory relationships. IReNA also provides functions to decode inferred regulatory networks, including the modularization of regulatory networks, the enrichment of transcription factors, and the construction of simplified regulatory networks among modules.

Several of these functions of IReNA are new, and not performed by other analytic tools. First, unlike current methods using scRNA-seq data for network analysis, IReNA integrates scRNA-seq data with scATAC-seq data to reconstruct regulatory networks. The results indicate that the integrated analysis of scRNA-seq and scATAC-seq data could more precisely identify known regulators than network analysis using scRNA-seq data alone. Second, IReNA could provide cell state-specific regulatory networks through modularizing regulatory networks. Regulatory networks are modularized according to the clustering results of gene expression profiles which represent different cell states and specific biological functions. Third, IReNA statistically analyzes modular regulatory networks and identifies reliable transcription factors including known regulators. Applied to multiple datasets, the method used in IReNA showed a consistently better performance on identification of known regulators than Rcistarget, which conducts transcription factor enrichment analysis based on the rank of all genes for each motif [13]. Fourth, we created a unique function in IReNA to construct simplified regulatory networks among modules which reveal key regulatory modules and factors, facilitating the interpretation of dynamic biological regulations.

IReNA can be used to infer regulatory networks for the same cell type, or for several cell types which are related by lineage during the development or disease progression. Correspondingly, cell type-specific or lineage-specific transcriptomes and epigenomes must be extracted prior to network inference. Transcriptomes could be measured by bulk RNA-seq or scRNA-seq, whereas epigenomes may be detected using either bulk ATAC-seq or scATAC-seq. Transcriptomes and epigenomes used in IReNA should be matched for the same cell type or the same condition. Currently, IReNA has used scRNA-seq and scATAC-seq data of unpaired cells from the same condition to perform network analysis. However, parallel scRNA-seq and scATAC-seq profiles from the same cells are emerging, and updates to IReNA will accommodate these new data.

In previous studies, we demonstrated that IReNA can be used to integrate scRNA-seq and ATAC-seq to reconstruct gene regulatory networks controlling retinal regeneration and retinal development [7,14]. Through network analysis using IReNA, we identified modular gene regulatory networks and key transcription factors regulating different cell states in retinal Müller glia in zebrafish and mice. Using genetic loss of function analysis, we confirmed that several transcription factors identified by IReNA are critical for retinal regeneration, including *hmga1a* and *yap1* in zebrafish and *Nfia/b/x* in mice [7]. These results indicate that IReNA can provide reliable regulatory networks and reveal key regulators during biological processes including retinal regeneration.

Using public scRNA-seq and ATAC-seq data from three studies, we further performed regulatory network analysis through IReNA, which provides meaningful biological insights. According to simplified intermodular regulatory networks in liver regeneration, organic acid catabolism-related modules and transcription factors are needed to be repressed to activate cell cycle progression in hepatocytes, e.g., hepatocyte nuclear factor 4 alpha (*Hnf4a*) (Figure 2F, 3C). The study in mouse liver regeneration demonstrated that deletion of *Hnf4a* leads to sustained proliferation [15]. In heart regeneration, simplified modular regulatory networks imply that regulatory modules and transcription factors controlling cardiac muscle contraction promote cell cycle of cardiomyocytes (Figure 4B). It has been reported that one of such factors *Gata4* could activate heart regeneration in zebrafish and mice [16,17]. In regulatory networks of NASH, we identified *Fos* and *Klf6*, which regulate TH17/Treg cells in NASH patients and are up-regulated in the context of steatohepatitis, respectively [18,19]. The results indicate that IReNA could be applied to identify key modules and transcription factors that regulate a range of different biological processes, although further functional validation of putative key regulators is required.

## Conclusion

In summary, we have developed IReNA to perform regulatory network analysis, including network inference, network modularization, transcription factor enrichment and construction of simplified regulatory networks among modules. IReNA showed a consistently better performance at identifying known regulators when integrating scRNA-seq data with scATAC-seq data to reconstruct gene regulatory networks. IReNA also outperformed the existing method Rcistarget on identifying known regulators. Key transcription factors and regulatory relationships identified by IReNA are potential targets controlling tissue regeneration, diseases, and other dynamic biological processes. Through the construction of modular regulatory networks and simplified regulatory networks among modules, IReNA facilitates the understanding of regulatory mechanisms and provides meaningful biological insights.

## Materials and methods

### Description of scRNA-seq, bulk ATAC-seq and scATAC-seq data

To demonstrate analysis flow of IReNA, we used public scRNA-seq and ATAC-seq data from three studies, which analyzed datasets obtained from models of liver regeneration, heart regeneration, and nonalcoholic steatohepatitis (NASH), respectively [11,20,21].

For liver regeneration, partial hepatectomy (PHx) was performed in adult mice [20]. The study conducted both scRNA-seq and scATAC-seq on liver tissues at 0 and 48 hours after PHx (accession number GSE158866 and GSE158873, available at Gene Expression Omnibus database https://www.ncbi.nlm.nih.gov/geo/). According to the original annotation of cell types in the study, there are 2815 and 7004 hepatocytes measured separately for scRNA-seq and scATAC-seq.

In the study of heart regeneration, scRNA-seq profiles were measured on 4884 cardiomyocytes at 1 day, 3 days after myocardial infarction, and 1 day after sham surgery in neonatal mice (accession number GSE130699) [22]. Meanwhile, scATAC-seq was performed on 755 cardiomyocytes at 3 days after myocardial infarction on postnatal day one (accession number GSE142365).

The scRNA-seq and bulk ATAC-seq data from the study of NASH are available through the accession numbers GSE128334 and GSE128335 [11]. In this study, scRNA-seq profiles were measured on 6184 non-parenchymal cells, including 2748 Kupffer cells, from liver tissues of healthy and NASH mice. Bulk ATAC-seq was conducted on Kupffer cells from two healthy and two NASH samples.

### Regulatory network inference

Prior to network inference, we identified differentially expressed genes (DEGs) and the expressed transcription factors as the potential genes in network analysis. Monocle (version 2.1.8) was used to analyze scRNA-seq data and to construct the trajectory [23]. Pseudotime of individual cells was further inferred and used to identify DEGs (q-value < 0.005, fraction of expressed cells > 10% and single-cell expression difference > 0.1). Single-cell expression difference was defined as previously described [7]. Given that some key transcription factors may not be DEGs, we included all expressed transcription factors which expressed in > 5% cells for network analysis.

According to the previous evaluation of the methods for network inference, we selected top performing method GENIE3 as the default method for network inference in IReNA [3,4]. GENIE3 infers regulatory relationships of transcription factors to target genes based on random forest regression [4]. GENIE3 (version1.16) was used to analyze scRNA-seq data and to calculate the weight of regulation for each gene pair. We selected gene pairs which have > 0.0001 weight. Meanwhile, only gene pairs which contain at least one transcription factor from the TRANSFAC database (version 2018.3) were chosen as potential regulatory relationships.

To determine activating and repressive regulatory relationships, we calculated Pearson’s correlation of gene pairs using smoothed expression profiles. When Pearson’s correlation > 0, the regulation type of gene pair is defined as activation. When Pearson’s correlation < 0, the regulation type is defined as repression.

### Identify transcription factor binding motifs in regulatory regions of targeted genes

If bulk or single-cell ATAC-seq data is not available, gene regulatory relationships inferred from scRNA-seq data analysis were further refined according to transcription factor binding motifs present in the promoter regions of genes. Fimo was used to identify transcription factor binding motifs in the promoter regions (ranging from 1000bp upstream to 500bp downstream of the transcription start sites) of the genes [24]. Position weight matrices of binding motifs were from TRANSFAC database (version 2018.3). Regulatory relationships were selected for further network analysis if the binding motif of transcription factor occurs in the promoter region of the target gene.

### Analyze bulk or single-cell ATAC-seq data to refine regulatory relationships

If bulk ATAC-seq data is available, we use the following six steps to preprocess raw data in fastq format. (I) Remove adaptors of pair-end raw reads using fastp software (version 0.21.0) [25]. (II) Align reads the GRCm38/mm10 genome using bowtie2 (version 2.4.1) with default parameters [26]. (III) Filter low-mapping-quality reads (MAPQ < 10) and exclude duplicated reads separately using Samtools (version 1.3.1) and Picard (http://broadinstitute.github.io/picard/) [27]. (IV) Call peaks through MACS2 (version 2.1.0) with the parameter extsize = 200 and shift = 100 [28]. (V) Use HTseq (version 0.12.4) to calculate the count number of each peak [29]. (VI) Combine the peaks across all samples to obtain the union peaks and identify differentially accessible peaks using EdgeR [30].

Different from bulk ATAC-seq data, scATAC-seq data was preprocessed through following steps. (I) Map raw sequencing data in fastq format to the reference genome (GRCm38/mm10) with cellranger (version 2.0.0) [31]. (II) Use ArchR (version 1.0.1) to integrate scATAC-seq and scRNA-seq data with unconstrained integration methods [12]. Identify peak-to-gene links by calculating the correlation between peak accessibility and gene expression across individual cells, and retain peak-to-gene links with absolute value of correlation > 0.2, FDR < 1E-6, varCutOffATAC (variance of peak accessibility) > 0.7 and varCutOffRNA (variance of gene expression) > 0.3.

After processing bulk or single-cell ATAC-seq data, the following steps were used to identify footprints and to refine regulatory relationships. (I) Identify the footprints of peaks through HINT (version 0.13.2) and select high-quality footprints (tag-count score > 80th percentile) for downstream analysis [32]. (II) Select footprints which are covered by differentially accessible peaks. (III) Run Fimo to find binding motifs in the footprints according to the position weight matrices of motifs from TRANSFAC database [24]. (IV) Identify footprint-related genes. For bulk ATAC-seq data, ChIPseeker (version 1.26.2) is used to annotate footprint regions. Genes related to footprint regions are considered as footprint-related genes [33]. For scATAC-seq data, genes linked by ArchR to peaks are considered as footprint-related genes. (V) Use Rsamtools (version 2.6.0) to obtain the sequencing depth of the mapped reads which is used to calculate the number of insertions at each position of footprints (https://bioconductor.org/packages/Rsamtools). (VI) Use the number of insertions to calculate footprint occupancy score (FOS), and then select regulatory relationships which have high FOS (FOS > 0.1) to reconstruct regulatory networks. FOS was calculated using the formula defined as previously described [7].

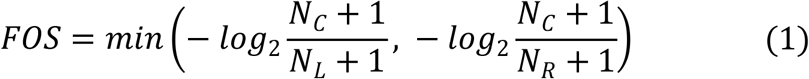

where *N*_*L*_, *N*_*C*_ and *N*_*R*_ are numbers of insertions separately in the left, center and right regions of the motif.

### Network modularization based on gene co-expression

Given the sparsity of scRNA-seq data, we used the smoothed expression profiles to perform gene co-expression analysis. The smoothed expression profiles were calculated according to pseudotime and branches on the trajectory. If there is a branch in the trajectory, pseudotime in each branch was divided into 20 equal intervals. Otherwise, pseudotime was divided into 50 equal intervals. Then, we calculated the average expression profile of single cells in each interval and obtained the smoothed expression profiles. DEGs and the expressed transcription factors were divided into different modules using the K-means clustering of the smoothed expression profiles. The optimal number of modules was determined by the silhouette coefficient calculated by R package ‘cluster’. Based on the modules of genes, the inferred regulatory networks were modularized. For each module, ClusterProfile (version 3.18.1) was used to perform functional enrichment analysis [34].

### Transcription factor enrichment for modular regulatory networks

Refined regulatory relationships were used to reconstruct modular regulatory networks of DEGs and expressed transcription factors. Cytoscape was used to display regulatory networks [35]. We performed the hypergeometric test to calculate the probability P that an individual transcription factor regulates a module, and then adjusted P values to false discovery rate (FDR) values. Enriched transcription factors which significantly regulate each module of genes were used to reconstruct regulatory networks. For the hypergeometric test, the probability was calculated as follows.

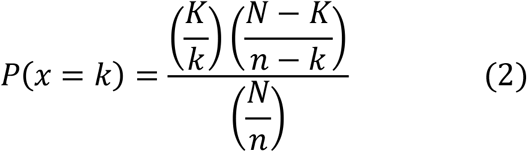

Here, 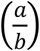 is a binomial coefficient. For identifying the significant transcription factor regulating module A, *N* and *n* represent numbers of all regulations and regulations targeting module A, respectively. *K* and *k* separately indicate the number of regulations from transcription factor and the number of regulations targeting module A from transcription factor.

### Construct simplified regulatory networks among modules

Based on modular regulatory networks of enriched transcription factors, we carried out another hypergeometric test to determine significant regulatory relationships among modules and reconstructed simplified regulatory networks among modules.

The same formula as (2) was used to calculate the probability of the regulation of module A to module B. In the formula, *N* and *n* represent numbers of all regulations and regulations from module A, respectively. *K* and *k* separately indicate the number of regulations from module B and the number of regulations from module A to module B.

The P values of regulations among modules were adjusted to FDR values. Regulatory relationships with FDR < 0.05 were regarded as significant regulations, and used to construct simplified regulatory networks among modules.

### Comparison of regulatory networks

To compare regulatory networks inferred from the integrated analysis of both scRNA-seq and ATAC-seq data, and from scRNA-seq data alone, we examined whether top and all enriched transcription factors in network analysis had been previously reported in the literature. Enriched transcription factors were ranked by FDR. For the study of nonalcoholic steatohepatitis, we used the biological terms ‘nonalcoholic steatohepatitis, steatosis or fatty liver disease’ to search the Google Scholar and PubMed databases. Given a limited number of studies reporting organ-specific regeneration, the common biological term ‘regeneration’ was used for each study of liver regeneration, and heart regeneration. We confirmed if the gene symbol and/or common gene name for individual transcription factors were present. If the gene symbol/gene name and biological term were both present in the title or the same sentence in the abstract, the biological function of the enriched transcription factor was regarded to have been reported in the literature.

To compare IReNA with existing methods for transcription factor enrichment, we used Rcistarget to identify key regulators analyzing the same scRNA-seq data [10]. Rcistarget calculates the normalized enrichment score (NES) which measures the enrichment of transcription factor binding motifs in the promoter regions of DEGs. We reconstructed regulatory networks for transcription factors which have > 3 NES. Then, we checked whether top (ranked by NES) and all transcription factors identified through Rcistarget were reported in the literature.

## Supplementary data

Supplementary data are available online.

## Acknowledgements

This work was supported by the National Natural Science Foundation of China [32170849], and the Guangdong Province Science and Technology Program [2020B1212060052].

## Availability of data and materials

IReNA is an open software and available online at https://github.com/jiang-junyao/IReNA.

## Author contributions

J.W., and J.Q. conceived the project. J.W., J.Q., and S.B. supervised the research. J.Y., P.L., and J.W. developed IReNA and performed sequencing data analysis to construct regulatory networks. J.Y., J.L., and S.H. checked transcription factors reported in the literature.

## Figure and table legends

**Figure S1.**
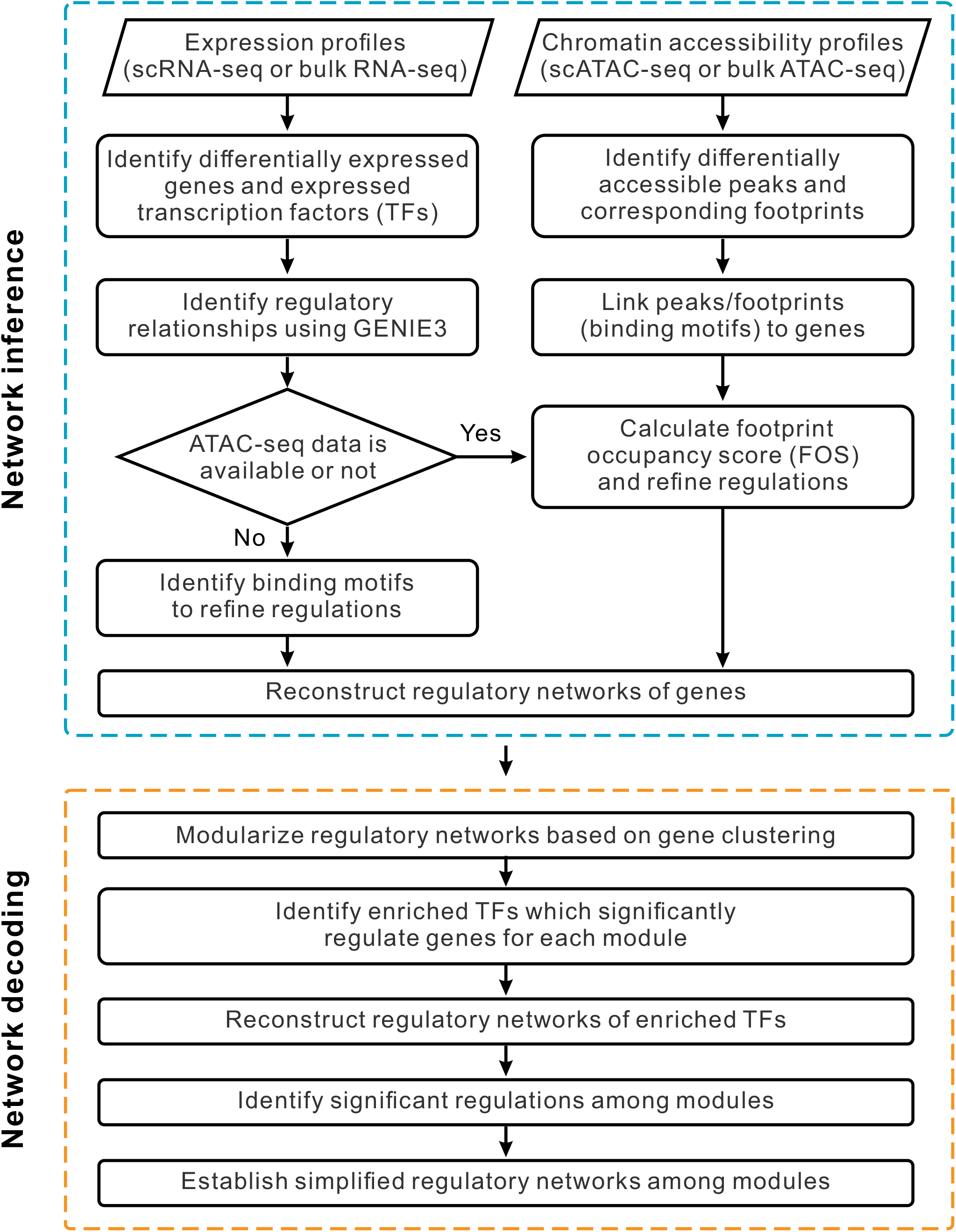
Workflow of IReNA. Differentially expressed genes (DEGs) and expressed transcription factors (TFs) are identified from single-cell or bulk RNA-seq expression profiles. Regulatory relationships among genes are inferred according to weights calculated by GENIE3. If ATAC-seq data is not available, binding motifs are further identified to refine regulatory relationships. Otherwise, both binding motifs and footprint occupancy score (FOS) are computed to refine regulatory relationships. If scATAC-seq is used, peaks are linked to genes through ArchR. After network inference, networks are decoded through modularizing regulatory networks, identifying enriched TFs, and establishing simplified regulatory networks among modules.

**Figure S2.**
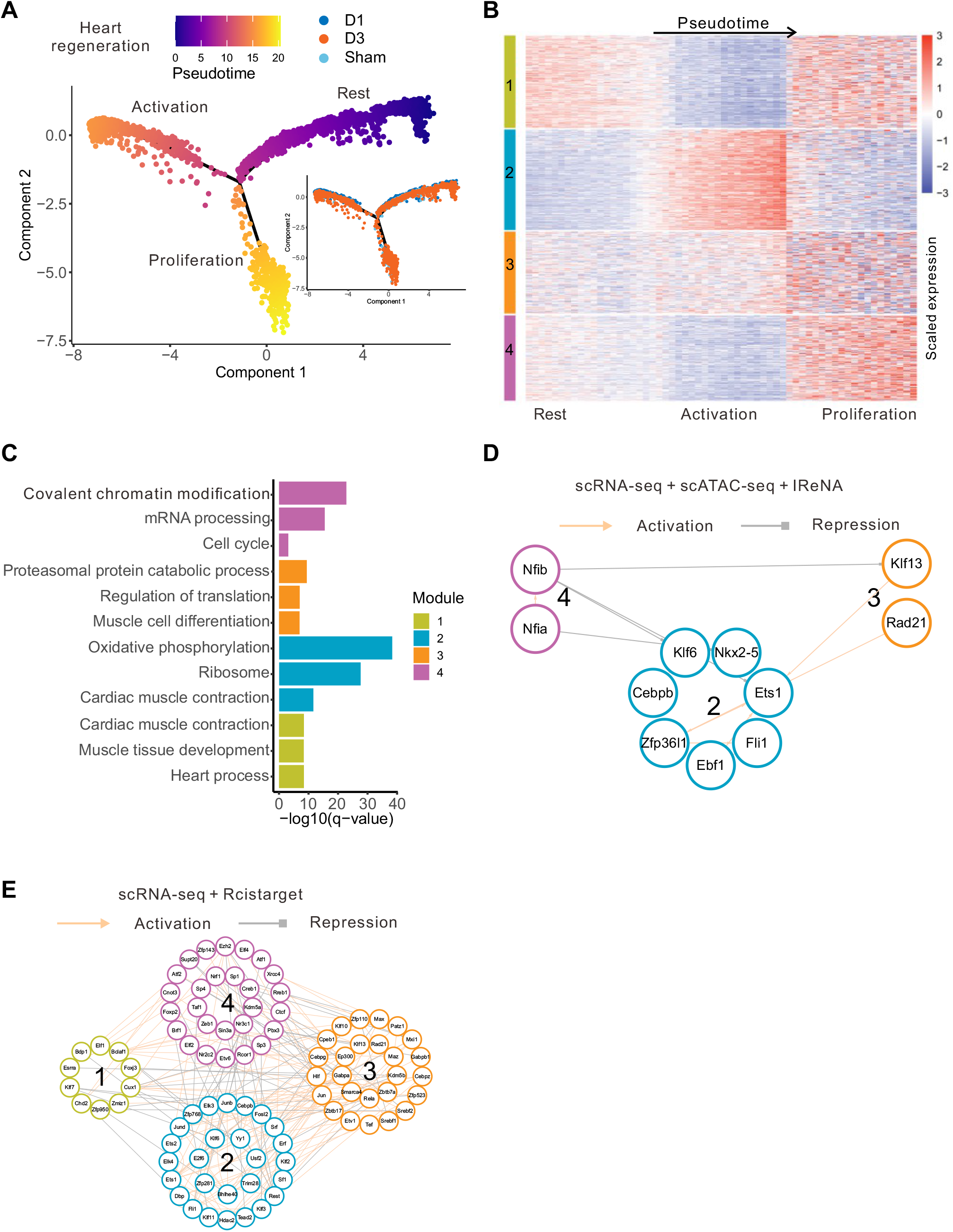
Regulatory network analysis of heart regeneration. (A) Trajectory of 4884 cardiomyocytes from neonatal heart tissues at day 1, day 3 after myocardial infarction, and day 1 after sham surgery. (B) 4340 DEGs and expressed transcription factors were divided into 4 modules according to K-means clustering of scRNA-seq expression profiles. Rows represent genes, and columns indicate intervals of pseudotime divided by branches. (C) Enriched functions of four modules of genes. Colors represent modules. (D) Regulatory networks of 11enriched transcription obtained through integrated analysis of scRNA-seq and scATAC-seq data. Grey edge represents negative regulation, and the yellow edge represents positive regulation. (E) Regulatory networks of 90 enriched transcription factors obtained by Rcistarget analysis of scRNA-seq data.

**Figure S3.**
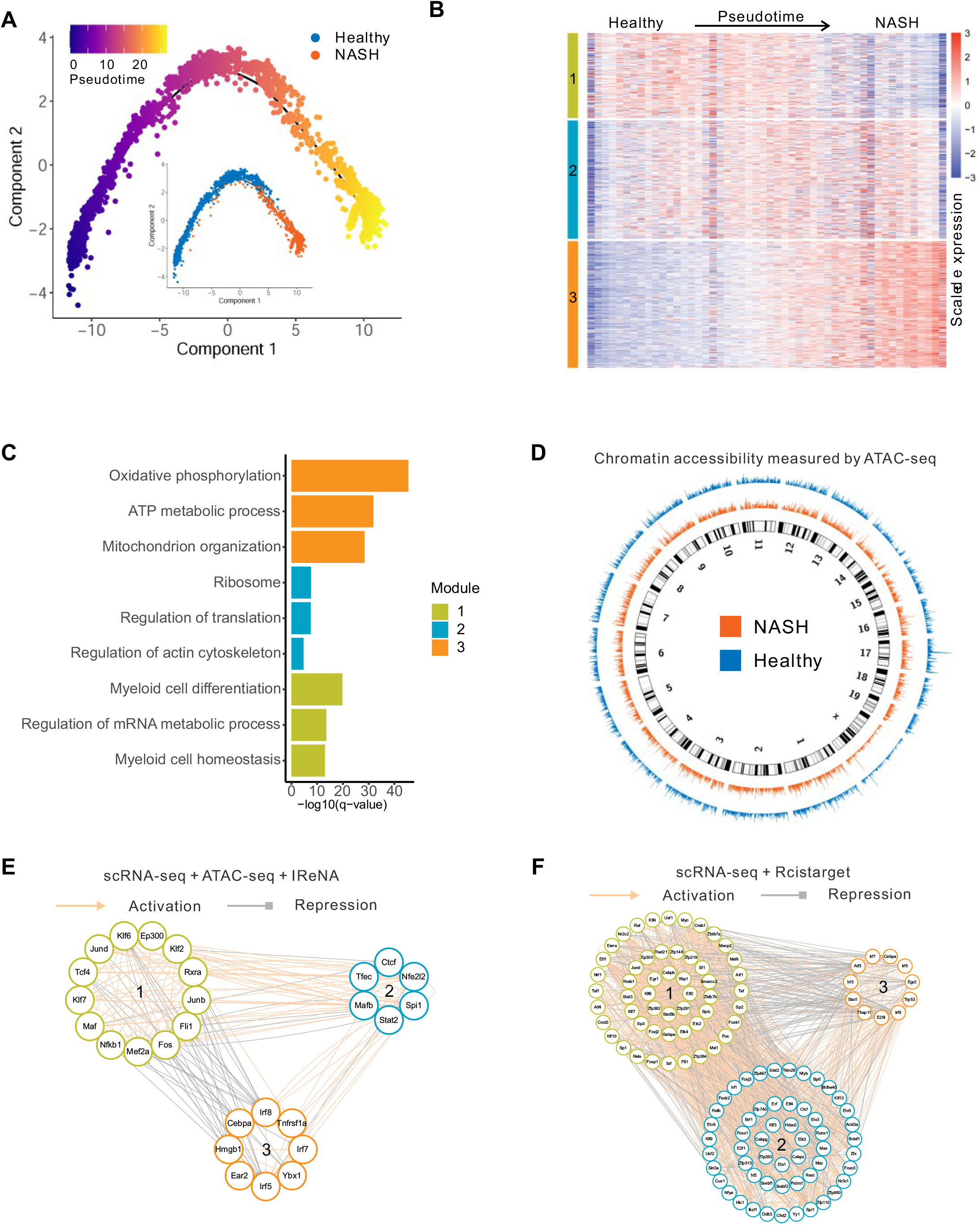
Regulatory network analysis of Kupffer cells from healthy and nonalcoholic steatohepatitis (NASH) liver tissues. (A) Trajectory of 2748 Kupffer cells separately colored by pseudotime and samples. (B) Heatmap of scRNA-seq expression profiles of 2742 differentially expressed genes and expressed transcription factors. K-means clustering was used to divide genes into three modules. Each row represents one gene, and each column indicates one interval of pseudotime. (C) Enriched functions of three modules of DEGs. (D) Genome-wide chromatin accessibility of healthy and NASH Kupffer cells measured by bulk ATAC-seq. (E) Regulatory networks of 27 enriched transcription factors obtained through integrated analysis of scRNA-seq and bulk ATAC-seq data. Color of each circle indicates the module. Grey edge represents negative regulation, and the yellow edge represents positive regulation. (F) Regulatory networks of 119 enriched transcription factors obtained by Rcistarget analysis of scRNA-seq data.

**Table S1**. The list of transcription factors enriched in liver regeneration, heart regeneration and nonalcoholic steatohepatitis. Transcription factors were enriched by IReNA analysis of scRNA-seq alone (scRNA-seq + IReNA), the integrated analysis of scRNA-seq and scATAC-seq data (scRNA-seq + scATAC-seq + IReNA) or Rcistarget analysis of scRNA-seq alone (scRNA-seq + Rcistarget). The related literature and sentences are also listed.

